# Methods to Utilize Pulse Wave Velocity to Measure Alterations in Cerebral and Cardiovascular Parameters

**DOI:** 10.1101/2023.06.22.546154

**Authors:** Andrea G. Marshall, Kit Neikirk, Bryanna Shao, Amber Crabtree, Zer Vue, Heather K. Beasley, Edgar Garza-Lopez, Estevão Scudese, Celestine N. Wanjalla, Annet Kirabo, Claude F Albritton, Sydney Jamison, Mert Demirci, Sandra A. Murray, Anthonya T. Cooper, George E Taffet, Antentor O. Hinton, Anilkumar K. Reddy

## Abstract

Alzheimer’s Disease (AD) is a global health issue, affecting over 6 million in the United States, with that number expected to increase as the aging population grows. As a neurodegenerative disorder that affects memory and cognitive functions, it is well established that AD is associated with cardiovascular risk factors beyond only cerebral decline. However, the study of cerebrovascular techniques for AD is still evolving. Here, we provide reproducible methods to measure impedance-based pulse wave velocity (PWV), a marker of arterial stiffness, in the systemic vascular (aortic PWV) and in the cerebral vascular (cerebral PWV) systems. Using aortic impedance and this relatively novel technique of cerebral impedance to comprehensively describe the systemic vascular and the cerebral vascular systems, we examined the sex-dependent differences in 5x transgenic mice (5XFAD) with AD under normal and high-fat diet, and in wild-type mice under a normal diet. Additionally, we validated our method for measuring cerebrovascular impedance in a model of induced stress in 5XFAD. Together, our results show that sex and diet differences in wildtype and 5XFAD mice account for very minimal differences in cerebral impedance. Interestingly, 5XFAD, and not wildtype, male mice on a chow diet show higher cerebral impedance, suggesting pathological differences. Opposingly, when we subjected 5XFAD mice to stress, we found that females showed elevated cerebral impedance. Using this validated method of measuring impedance-based aortic and cerebral PWV, future research may explore the effects of modifying factors including age, chronic diet, and acute stress, which may mediate cardiovascular risk in AD.

**New and Noteworthy:** Here, we presented a new technique which is an application of the concept of aortic impedance to determining cerebral impedance. While aortic PWV is typically utilized to study aortic stiffness, we also developed a technique of cerebral PWV to study cerebral vascular stiffness. This method may be useful in improving the rigor of studies that seek to have a dual focus on cardiovascular and cerebral function.

## Introduction

Alzheimer’s disease (AD) is a progressive neurodegenerative disorder that affects millions of people worldwide, a rate that is expected to grow with the aging population (1). AD causes deficits in quality of life and daily living marked by a decline of memory and cognitive functions (2). A hallmark feature of AD is the accumulation of aggregated amyloid beta (Aβ) protein in the brain, which ultimately leads to synaptic dysfunction, neuronal loss, and cognitive impairment (3). In addition to genetic factors, lifestyle and environmental factors such as stress and diet have been implicated in the development and progression of AD (4). Beyond dementia, AD also impacts cardiovascular function, as the amyloid beta plaques also accumulate in the heart (1). A key risk factor of AD is hypertension, and it has been hypothesized that neuroinflammatory effects link these pathologies (5). However, the full extent of the mechanisms which link AD and cardiovascular disease remains poorly understood. Here, we sought to expand the understanding of how AD alters cardiovascular and cerebral blood flow in conditions of chronic diet changes or acute stress.

AD is well understood to cause neuropsychologic decline but is also understood to affect cardiovascular dynamics. AD importantly is shown to potentially be associated with hypertension (HTN) through the formation of neuritic plaques in cerebral vasculature and potentially limiting cerebral blood flow (6). In human populations, even with correcting for risk factors such as age, HTN reduction is correlated with increased retention of cognitive ability, indicating blood pressure treatment as a potential mechanism to treat AD (1). This blood pressure linkage to AD is especially pronounced when it comes to the systolic pressure (7). However, a past review on the matter shows the issue to be far more complicated, as many previous studies are not using standardized HTN measurements and there may be age-dependent and ethnicity-dependent influences of HTN on AD development (8). Together, this underscores the need to better understand hemodynamics in AD.

Hemodynamics that includes pulse wave velocity (PWV) is important to study aortic/arterial stiffness and cardiovascular function (9). Even when adjusting for factors including age, race, and disease states, National Health and Nutrition Examination Survey studies indicate PWV remains a strong predictor of overall mortality in human populations (10). Past studies looking at echocardiographic parameters in humans with AD showed that diastolic function was more heavily impacted in AD pathology, marked by increased arterial stiffness, increased atrial conduction times, reduced blood flow, and altered mitral valve velocities (11). Carotid-femoral pulse wave velocity increased aortic stiffness has also been linked as an independent predictor of cognitive impairment for both dementia and AD (12). Another longitudinal study across over 1700 participants found that arterial stiffness and alterations in pulse pressure are antecedents for cognitive decline (13). PVW can further be used to distinguish vascular dementia and AD, with the former often expressing increased vascular stiffness relative to AD (14). Past studies have also highlighted the importance of looking at both PWV as well as cerebral blood flow, through MRI, with hypertensive symptoms altering heart and brain structure (15). Yet there remains controversy, as some studies have found no correlation between carotid-femoral PWV and AD (16).

Our primary way to study AD cardiovascular-related parameters is through 5x transgenic (5xFAD) mice, which have 5 AD-linked mutations. The 5xFAD mouse model expresses human amyloid precursor protein (APP) and presenilin 1 (PS1) mutations, leading to the rapid accumulation of Aβ and the development of AD-like pathology (17). This model also bears numerous similarities to human models including Aβ-butyrylcholinesterase association and hallmarks of progressive loss of cognitive function concomitant with reduced synaptic markers (18). Due to apoptotic neuron loss, 5xFAD mice display memory deficits by 4 months of age (19). This mouse model has been widely used to study the mechanisms underlying AD and to test potential therapeutic interventions. Past studies using 3x transgenic mice models highlight that with induced hypertension, mice had faster development of AD marked by upticks in Aβ, amyloid plaque load, and phosphorylated tau (20). This validates that transgenic mice models retain the relationship between cerebrovascular and cardiovascular dysfunction that has been observed in AD.

To advance a new way of studying cerebrovascular dynamics in both wildtype and 5XFAD mice, we developed a technique to measure cerebral pulse wave velocity. The relationship between PWV and AD pathology in 5xFAD models remains unclear, especially as it comes to cerebral pulse wave velocity (cPWV), a hemodynamic parameter that has been increasingly recognized as an important predictor of cerebrovascular disease. While aortic stiffness can serve as a predictor for strokes, it is limited in detection of minor changes which precede cerebrovascular changes such as microbleeds (21). Similar to its aortic counterpart, cPWV measures the speed at which arterial pressure or velocity waves propagate through the cerebral arteries. Measuring cerebral impedance is a novel measure of the comprehensive characterization of cerebral blood vessels, which is highly relevant as the brain is a highly vascularized organ that requires a constant supply of oxygen and nutrients, with disruptions in blood flow being linked to cognitive impairment. In AD, this remains highly relevant as it may relate to Aβ clearance (8). A recent clinical trial protocol has proposed utilizing carotid-cerebral PWV as a mechanism to better understand acute ischemic strokes (22). Here, we use pulsed Doppler ultrasound to simultaneously measure the arrival of the velocity wave at two different arterial sites, one at the aortic arch and another at ophthalmic artery in the distal internal carotid artery and measure the physical distance between the two sites to estimate cerebral PWV (cPWV). Also, using carotid flow velocity and aortic blood pressure we measured cerebral impedance and impedance-based PWV (cPWV_Zc_) which is crucial for improving our understanding of the pathophysiology of AD (Table 1).

To validate our methods, we studied sex-dependent, diet-dependent, and stress-induced differences in WT and 5XFAD mice. While it is clear that females are more likely to get AD in humans, there remains a gap in the literature in understanding sex-dependent differences in cerebral hemodynamics in 5xFAD mouse models (18). Using a combination of aortic and cerebral PWV, our findings provide important insights into the relationship between PWV and AD pathology, as well as potential novel mechanisms of protection. By understanding how these factors interact to influence arterial stiffness and cognitive function, we may be able to identify new therapeutic targets for the treatment and prevention of AD.

## Methods Before You Begin

### Animals

All animal protocols were approved by the Institutional Animal Care and Use Committee of Baylor College of Medicine in accordance with the National Institutes of Health Guide for the Care and Use of Laboratory Animals. We used 3 groups of 5xFAD mice at 4-5 months of age. The diets of the 3 groups of mice consisted of standard commercial chow (2920X Harlan Teklad, Indianapolis, IN, USA), high-fat diet (60% kCal from fat), and Equi diet () with free access to food and water (**Figure S1**). Separately, we also used 8-9 month C57BL6J mice with chow diet only. All the mouse groups are shown below.

| Animals | Strain | Male | Female | Age | Diet |
| --- | --- | --- | --- | --- | --- |
| Group1 | 5xFAD | n=10 | n=9 | 4-5mo | Normal Chow |
| Group2 | 5xFAD | n=10 | n=9 | 4-5mo | HFD |
| Group3 | 5xFAD | n=5 | n=5 | 4-5mo | Equi Diet (Pre & Post Stress) |
| Group4 | C57BL6/J | n=5 | n=5 | 8-9mo | Normal Chow |

Mice were initially anesthetized with 2.5% isoflurane in the induction chamber and then transferred to a heated (37±1°C) electrocardiography (ECG) board (MouseMonitor S, Indus Instruments, Webster, TX) with the paws taped to the ECG electrodes and isoflurane maintained at 1.5% via nose cone.

### Doppler flow velocity measurements

We used a 20 MHz Doppler probe to measure doppler aortic aortic arch velocity, ophthalmic artery velocity (representative of cerebral blood flow), and abdominal aortic velocity signals to determine cerebral PWV (cPWV) and aortic PWV (aPWV) (figure 1a). We also measured aortic outflow velocity and carotid flow velocity along with aortic blood pressure, and mitral blood flow velocity (figure 1b) to determine cardiac and cerebral hemodynamics. All signals were acquired and stored using Doppler Flow Velocity System (DFVS; Indus Instruments, Webster, TX). We measured separation distance between aortic arch and OA sites and between aortic arch and abdominal aortic sites to determine cPWV and aPWV, respectively. We measured peak and mean aortic velocities, stroke distance (Sd), aortic ejection time (ET), peak and mean aortic accelerations from aortic outflow velocity, and early peak and atrial peak velocities, E/A ratio, E deceleration time, isovolumic contraction (IVCT) & relaxation (IVRT) times from mitral inflow signals, and myocardial performance index (also known as Tei index = (IVCT + IVRT)/ET). From the carotid flow velocity signal, we calculated peak, minimum, and mean velocities, pulsatility index, and resistivity index are calculated.

**Figure 1:**
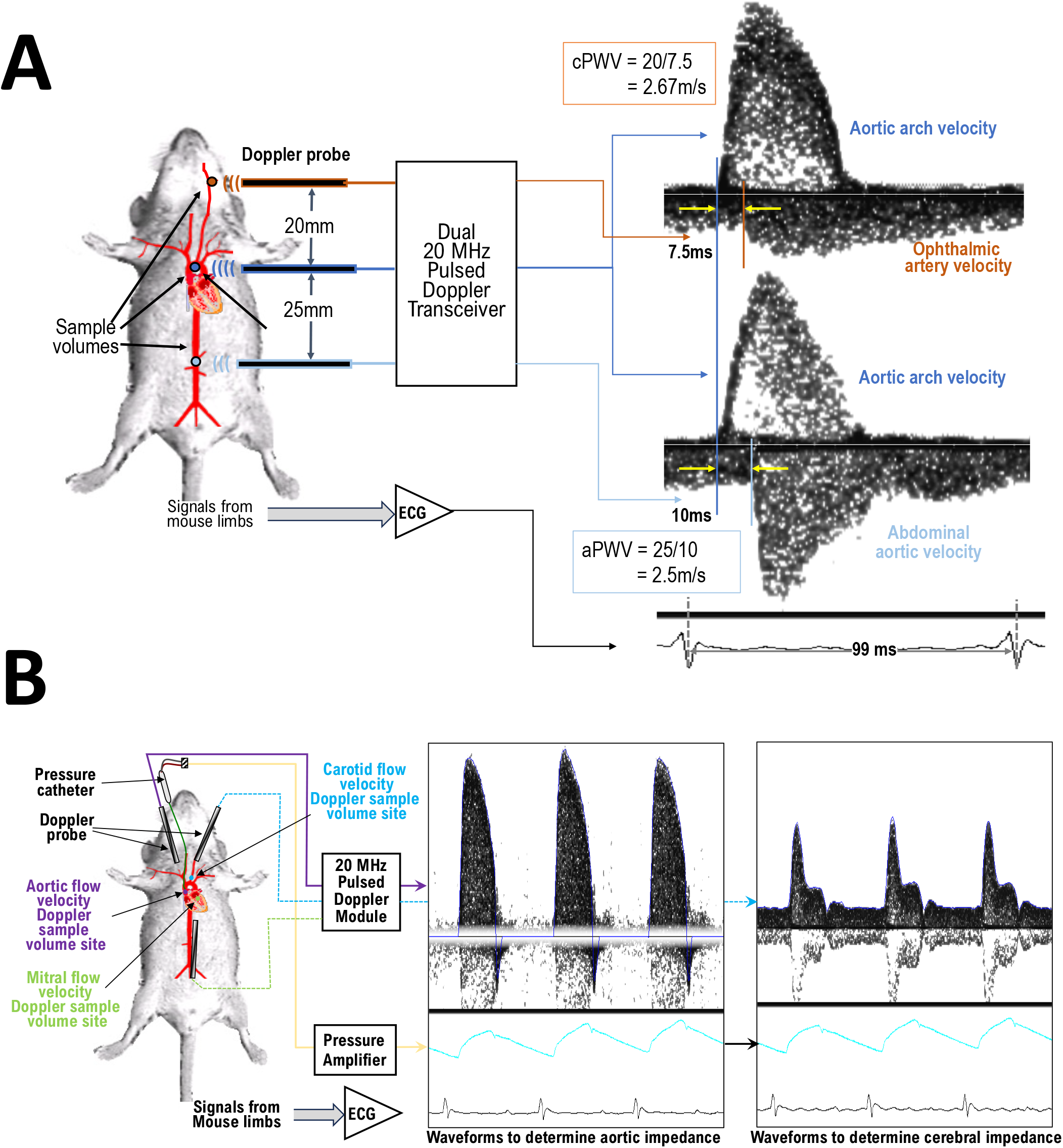
**(A)** Experimental setup to noninvasively measure cerebral (at ophthalmic artery, OA) & aortic pulse wave velocity. One probe is fixed at the aortic arch and the other probe is switched between ophthalmic artery or abdominal aorta sites for Arch-OA combined signals or Arch-Abd combined signals. Electrocardiography (ECG) waveform is shown for timing. (**B**) Experimental setup to measure aortic blood flow velocity, aortic blood pressure and) ECGin mice. The probe is repositioned at carotid artery (shown in the right panel along with pressure & ECG signals). The probe is repositioned again to measure mitral flow velocity (waveform not shown).

### Blood pressure measurements

Blood pressure measurements were made as previously described (23–25), Briefly, a 1-French (0.33mm diameter) blood pressure catheter (SPR-1,000: Millar Instruments, Inc., Houston, TX) was introduced via the isolated right carotid artery and advanced into the ascending aorta to measure aortic pressure. About 2-3 second segments of blood pressure signals were acquired (simultaneously with either aortic flow velocity or carotid flow velocity and ECG signals) with the DFVS system. Systolic (SBP), diastolic (DBP), mean (MBP), pulse pressures (PP), end-systolic pressure (ESP) and rate-pressure product (RPP) were calculated from the recorded aortic blood pressure signals.

### Determination of aortic and carotid impedance

The method to determine aortic impedance was described elsewhere (22, 24-26). Aortic impedance is determined using aortic pressure-velocity relationship (figure 2a). The foot of aortic pressure waveform was aligned with the foot of the aortic velocity waveform to avoid potential errors in phase relation between pressure and velocity signals. The signals are converted to frequency domain using fast Fourier transform and impedance (|*Z*| |*P*|*/*|*V*|) parameters (peripheral vascular resistance [a*Z*_0_], characteristic impedance [a*Z*_C_], and impedance at first harmonic [a*Z*_1_]) are calculated. Aortic pulse wave velocity was calculated as aZ_C_/ρ (ρ-density of blood). The foot of the blood pressure waveform was aligned with the foot of carotid flow velocity waveform and cerebral impedance was calculated in the same way as aortic impedance (figure 2b) and the cerebral impedance parameters (cerebral vascular resistance [c*Z*_0_], cerebral characteristic impedance [c*Z*_C_], and cerebral impedance at first harmonic [c*Z*_1_]) were calculated. Cerebral pulse wave velocity was calculated as cZ_C_/ρ (ρ-density of blood).

**Figure 2:**
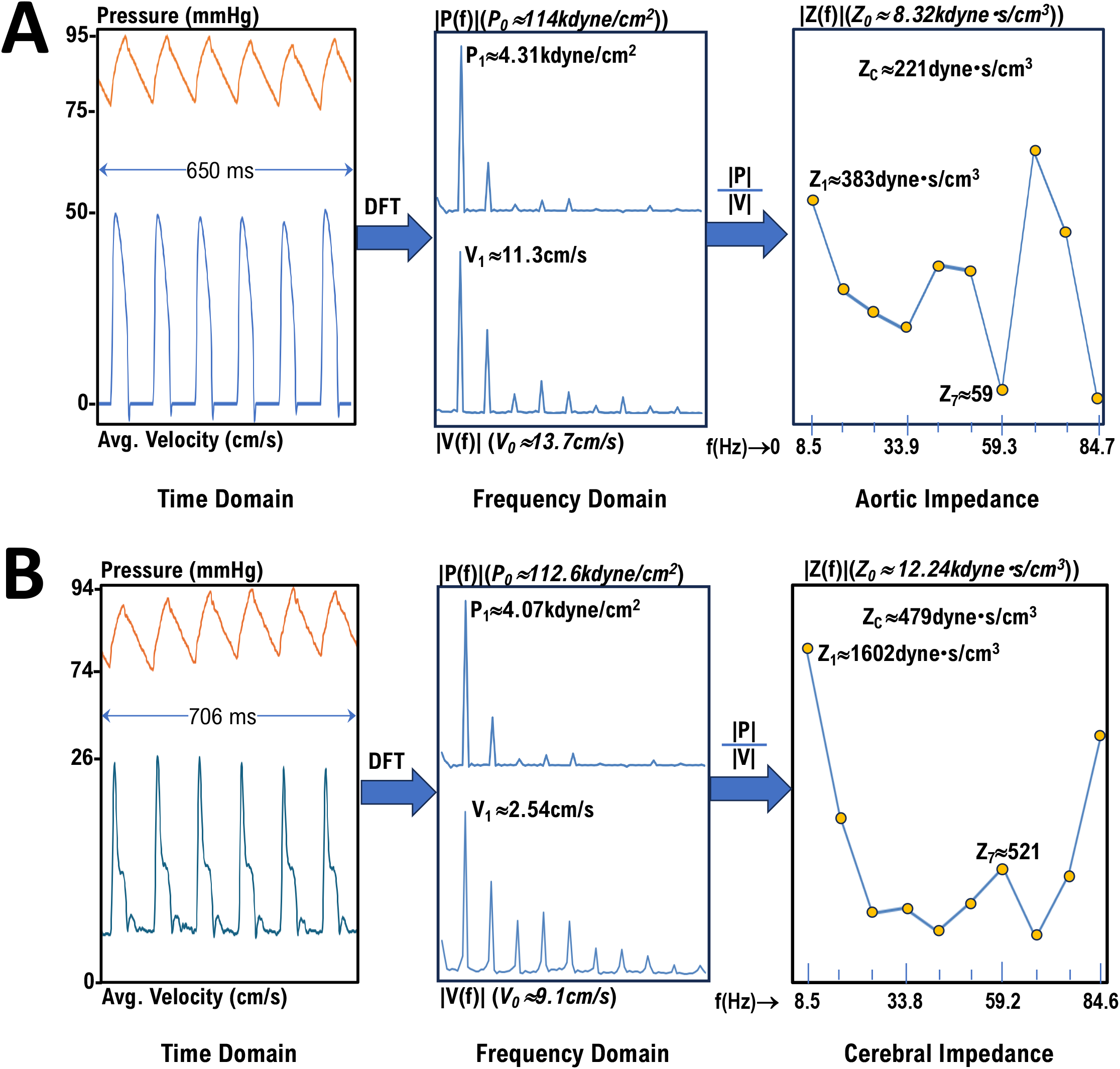
**(A)** Procedure to calculate aortic impedance – conversion of time domain signals to frequency domain spectrums and calculation of impedance modulus (|Z(f)| = |P(f)|/|V(f)|). Z_c_ is calculated as the average of Z_2_ to Z_10_ harmonics. **(B)** Procedure to calculate cerebral impedance – conversion of time domain signals to frequency domain spectrums and calculation of impedance modulus (|Z(f)| = |P(f)|/|V(f)|). Z_c_ is calculated as the average of Z_2_ to Z_10_ harmonics.

### Calculation of parameters to determine VVC

Elastance was determined as previously discussed (25, 26). Arterial elastance (Ea) was calculated as ESP/SV (stroke volume, SV = Sd * aortic cross-sectional area), end-systolic elastance (Ees) was calculated as ESP/ESV, ventricular-vascular coupling (VVC) was calculated as Ea/Ees, and stroke work (SW) was calculated as ESP*SV (27).

### Statistical analyses

All the data are presented as mean ± standard error of the mean (SEM). Dots represent each sample, as sample size varies throughout. Statistical analyses were performed via analysis using an unpaired T-test to compare conditions in each sex through Prism (GraphPad Software; La Jolla, USA).

### Step-By-Step to Measure Aortic and Carotid Velocity and Blood Pressure to Determine Aortic and Cerebral Impedance

#### Non-invasive

*<u>NOTE</u>*: Only allows for PWV measurement.

<u>Before you begin</u>: This protocol needs a Doppler system (see **Figure S2** – which shows the workflow of signals, but it does not require the pressure system for the noninvasive measurement of PWV).

1. On the day of the study, weigh the animal and anesthetize it in the induction chamber using 3.0% isoflurane (mixed with 1L/min 100% O2).
2. Transfer the animal to the heated ECG board, and place in supine position with 1.5% isoflurane supplied via a nose cone.
3. Apply artificial tears lubricant gel to the eyes to prevent dryness.
4. Apply ECG cream to the four paws and tape them to the ECG electrodes. *<u>NOTE</u>*: Ensure excess gas is scavenged for the safety of the operator.
5. Remove the hair from a small area near sternal border and a small area near mid abdominal area. *<u>NOTE:</u>* The temporal canthus site near the eye used for the internal carotid artery branch of the ophthalmic artery does not need hair removal.
6. Using a 20 MHz pulsed Doppler probe, place the tip on the chest at the aortic arch site and aim toward mid-line to find an aortic arch signal. Place the probe in a tightly held holder after the signal is found.
7. Holding the second probe by hand, place its tip at the temporal canthus of the mouse eye and aim the probe toward the internal carotid artery to measure ophthalmic artery (OA) flow velocity. Measure the separation distance between the two sites (where Doppler tips are placed).
8. Treat the Arch & OA signals as I & Q (**Figure S1**) and combine to produce signals (**Figure 1A**); save a 2-3 second segment of these signals. Determine the pulse transit time offline and calculate cPWV (see upper right-hand panel **Figure 1A**).
9. Move the second probe to the abdominal location to find the abdominal aortic signal. Measure the separation distance between the two sites.
10. Treat the Arch & Abd signals as I & Q and combine them to produce signals. Again, save a 2-3 second segment of these signals, determine pulse transit time offline, and calculate aPWV (see lower right panel in **Figure 1A**).
11. Once the measurements are made, wake the mouse up and return to the cage.

### Invasive

#### Before you begin

This specific protocol needs a Doppler system (see **Figure S2**-which shows the workflow of signals), that allows for the measurement and acquisition of Doppler velocity signals and ECG and blood pressure signals, simultaneously. A Millar pressure catheter and amplifier system are needed to measure blood pressure signals. The Doppler velocity signal and blood pressure are acquired simultaneously along with ECG for the temporal alignment of signals.

*<u>NOTE</u>*: Avoid the usage of analgesics as this may suppress blood pressure.

1. On the day of the study, weigh the animal and anesthetize in the induction chamber using 3.0% isoflurane (mixed with 1L/min 100% O2).
2. Transfer to heated ECG board and place in supine position with 1.5% isoflurane supplied via nose cone.
3. Apply artificial tears lubricant gel to eyes to prevent dryness.
4. Apply ECG cream to the four paws and tape them to the ECG electrodes. *<u>NOTE</u>*: Ensure excess gas is scavenged for the safety of the operator.
5. Shave the hair from the neck area and apply hair removal cream to ensure all body hair is removed.
6. Perform a pinch test to make sure the animal is unresponsive. If responsive, adjust isoflurane level to 2.0% and then return to 1.5% and redo the pinch test.
7. Use a rectal temperature probe to monitor body temperature and adjust the heated board to maintain body temperature at 37.0±0.5 °C. *<u>NOTE</u>*: The heated board temperature is usually higher than the body temperature, so this should be closely monitored.
8. Make a 60-70mm cut in the skin of the neck, to the right of the midline.
9. Expose the right carotid artery and cannulate with a pre-calibrated saline-soaked 1F Millar catheter and secure with a suture.
10. Advance the catheter advanced into the aorta. *<u>NOTE</u>*: Make sure the open skin is covered with wet gauze. Waveforms: (See **Figure 1B** for devices/probe placements for respective measurements)
11. Using the Millar catheter, measure aortic blood pressure signals continuously.
12. Aim a 20 MHz pulsed Doppler probe tip at right suprasternal notch, caudally towards the heart with a low angle for the measurements of aortic blood flow velocity.
13. For the measurement of carotid blood flow velocity, place the Doppler probe tip left of the midline in the neck and aim caudally towards the heart at as low an angle as possible.
14. For the measurement of mitral flow velocity, reposition the Doppler probe tip under the xiphoid and aim rostrally toward the heart.
15. Record Doppler signals. Doppler signals are processed in real time and displayed as Doppler spectrograms on the screen along with the blood pressure and ECG waveforms.
16. Record the following sets of signals/waveforms: **a**. Aortic blood pressure & flow velocity and ECG; **b**. Aortic blood pressure & carotid flow velocity and ECG; and **c**. Mitral flow velocity signals with ECG. *<u>NOTE</u>*: For each of the signal sets, collect 2-3 second segments of data for offline analysis.
17. Extract aortic and carotid flow velocity waveforms along with respective blood pressure waveforms.
18. Calculate aortic impedance and cerebral impedance, as shown in **Figures 2A & 2B**, respectively. Using a similar method to how aortic impedance is traditionally measured, cerebral impedance is derived from pressure waveforms of carotid velocity (**Table 1**). Use aortic pressure and carotid signal to equalize these values. *<u>NOTE</u>*: Discrete Fourier transform can be computed using a simple custom code, MATLAB, or other programs.

## Representative Results

While we measured general parameters, such as body weight and aortic cross-sectional area, the aim of this study was to validate our technique across experimental conditions (**Figure S1**). To begin with, we looked at wildtype mice with normal chow diet at an age of 8 months old, representing a standard model. Sex-dependent differences in LV afterload were evaluated using aortic impedance, which showed non-statistically significant differences of impedance parameters Zp, Z1, Zc, or PWVz (**Figure 3A-D**). When performing the cerebral counterpart to this measurement, while many measurements remained non-significant, females exhibited a lower cerebral Zp (**Figure 3E-H**).

**Figure 3:**
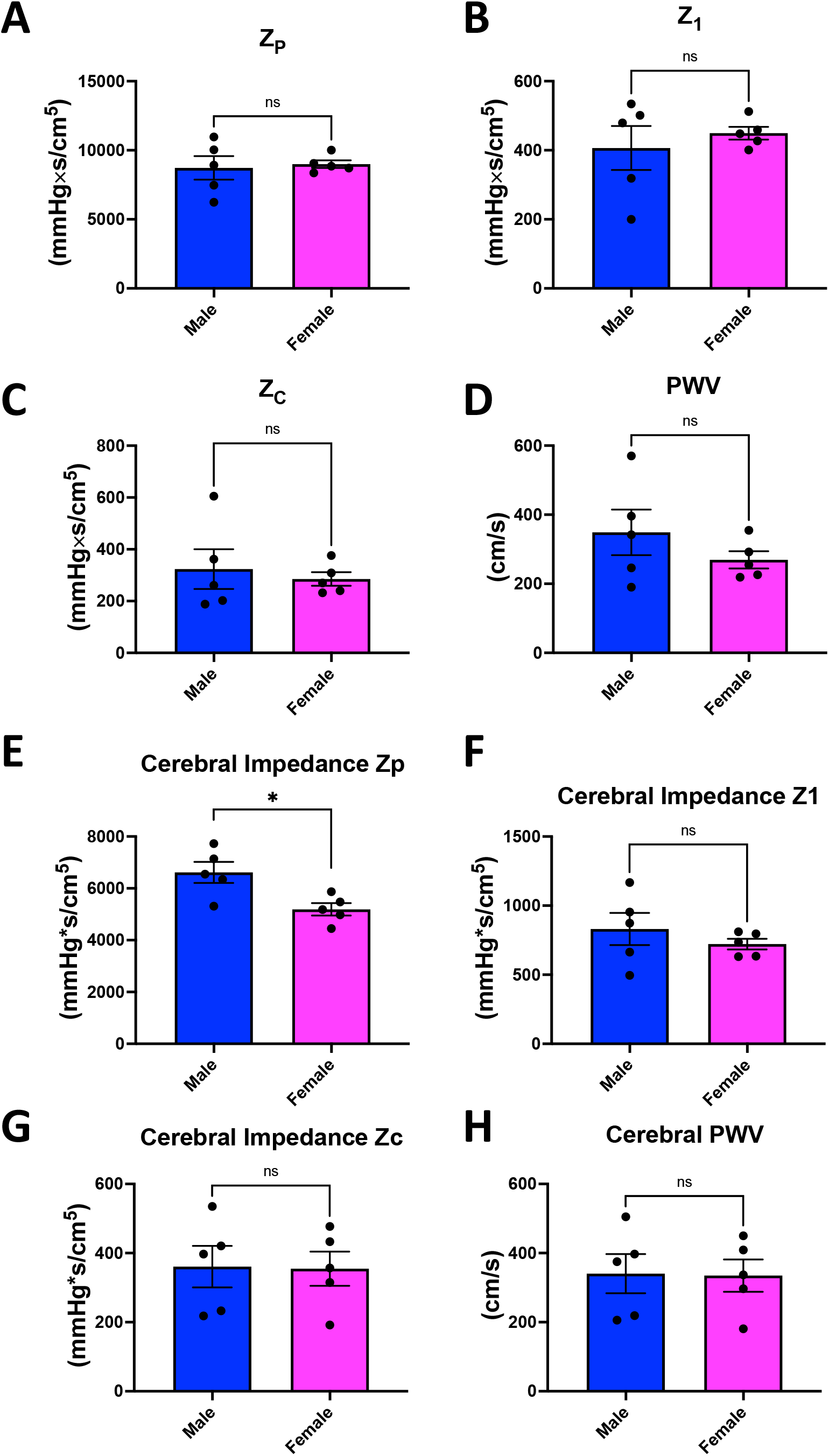
Parameters of aortic and cerebral impedance sex-dependent differences in chow-fed 8-month-old wildtype (C57BL6J) mice. (**A**) Total peripheral resistance (Z_P_), (**B**) impedance at first harmonic (Z_1_), (**C**) characteristic impedance, (Z_C_) (**D**) and impedance-based pulse wave velocity (PWV) in aortic and (**E-H**) cerebral impedance. Data are presented as mean±SEM (n = 5/group). * represents p <0.05, ns indicates a statistically non-significant relationship, as determined through an unpaired t-test.

To recapitulate these findings in a transgenic model, we looked at sex-dependent differences in aortic and cerebral impedance in 5XFAD mice chow-fed cohort that was 5-months old. LV afterload was evaluated using aortic impedance, which showed non-statistically significant differences in impedance parameters Zp, Z1, Zc, or PWVz (**Figure 4A-D**), but slightly more intracohort variability. When examining cerebral PWV, we found that both Zc and PWVz were both significantly lower in female mice (**Figure 4E-H**). These findings indicate that fine differences at baseline conditions may be detected, as well as showing that the 5XFAD mice differed in sex-dependent differences, as compared to WT mice. From there, we sought to validate the applicability of this aortic and cerebral measurement method with changes in diets.

**Figure 4:**
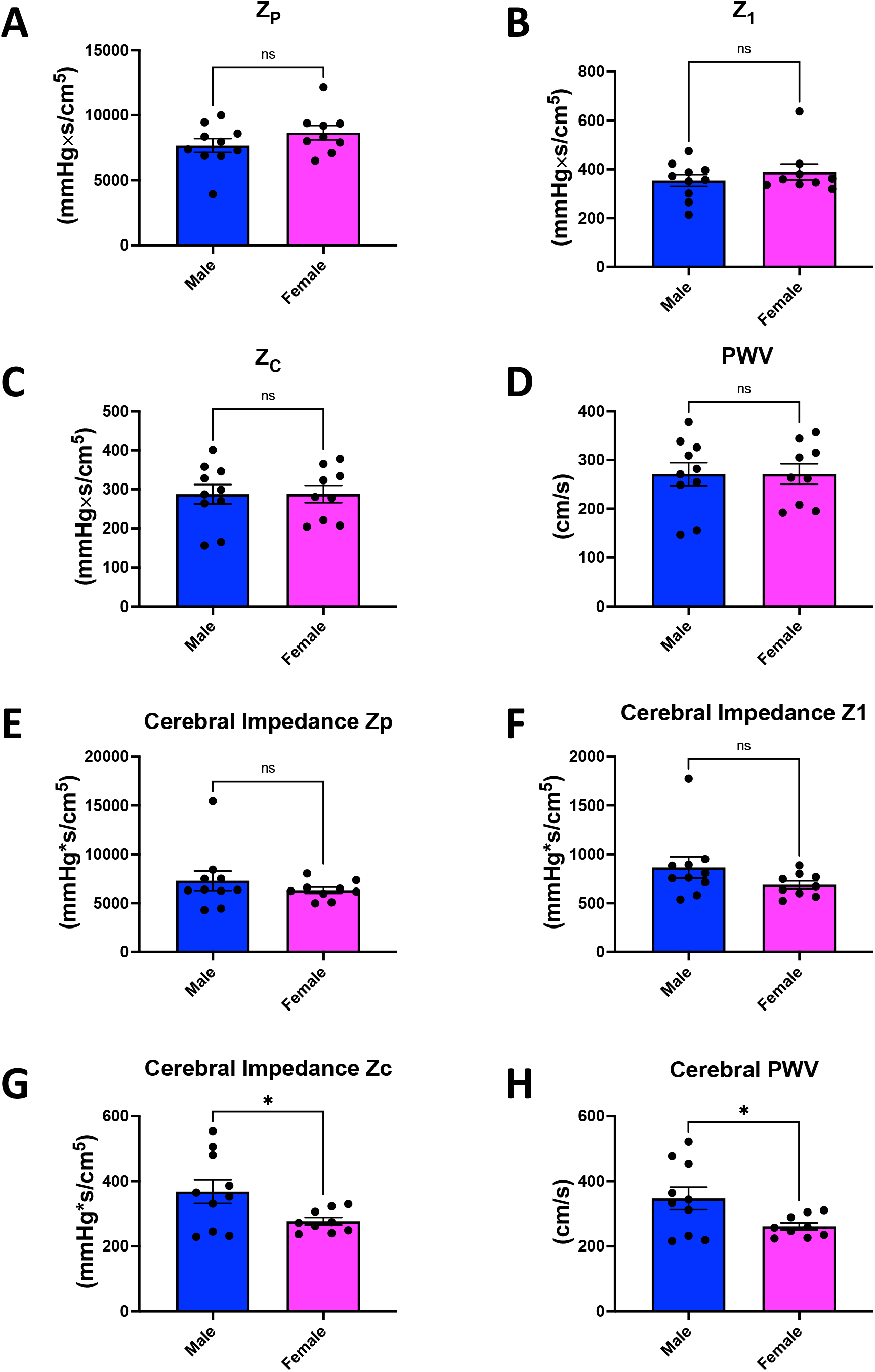
Parameters of aortic and cerebral impedance sex-dependent differences in chow-fed 5-month-old 5XFAD mice. (**A**) Total peripheral resistance (Z_P_), (**B**) impedance at first harmonic (Z_1_), (**C**) characteristic impedance, (Z_C_) (**D**) and impedance-based pulse wave velocity (PWV) in aortic and (**E-H**) cerebral impedance. Data are presented as mean±SEM (n = 9-10/group). * represents p <0.05, ns indicates a statistically non-significant relationship, as determined through an unpaired t-test.

A cohort of 8-week-old 5XFAD mice was subject to an HFD for 12 weeks and compared to the aforementioned cohort of 5XFAD mice on a chow diet. Generally, as compared to the chow cohort, the HFD diet cohort did not exhibit a significant difference in aortic or cerebral PWV parameters (**Figure S3**). However, when looking at sex-dependent differences within the 5XFAD cohort fed an HFD, we observed that cerebral parameters no longer showed a significant difference, but aortic Zc and PWVz were significantly lower in female mice (**Figure 5A-H**). These results that our aortic and cerebral methods have the sensitivity to detect minute sex-dependent differences that arise as a result of changes in diet.

**Figure 5:**
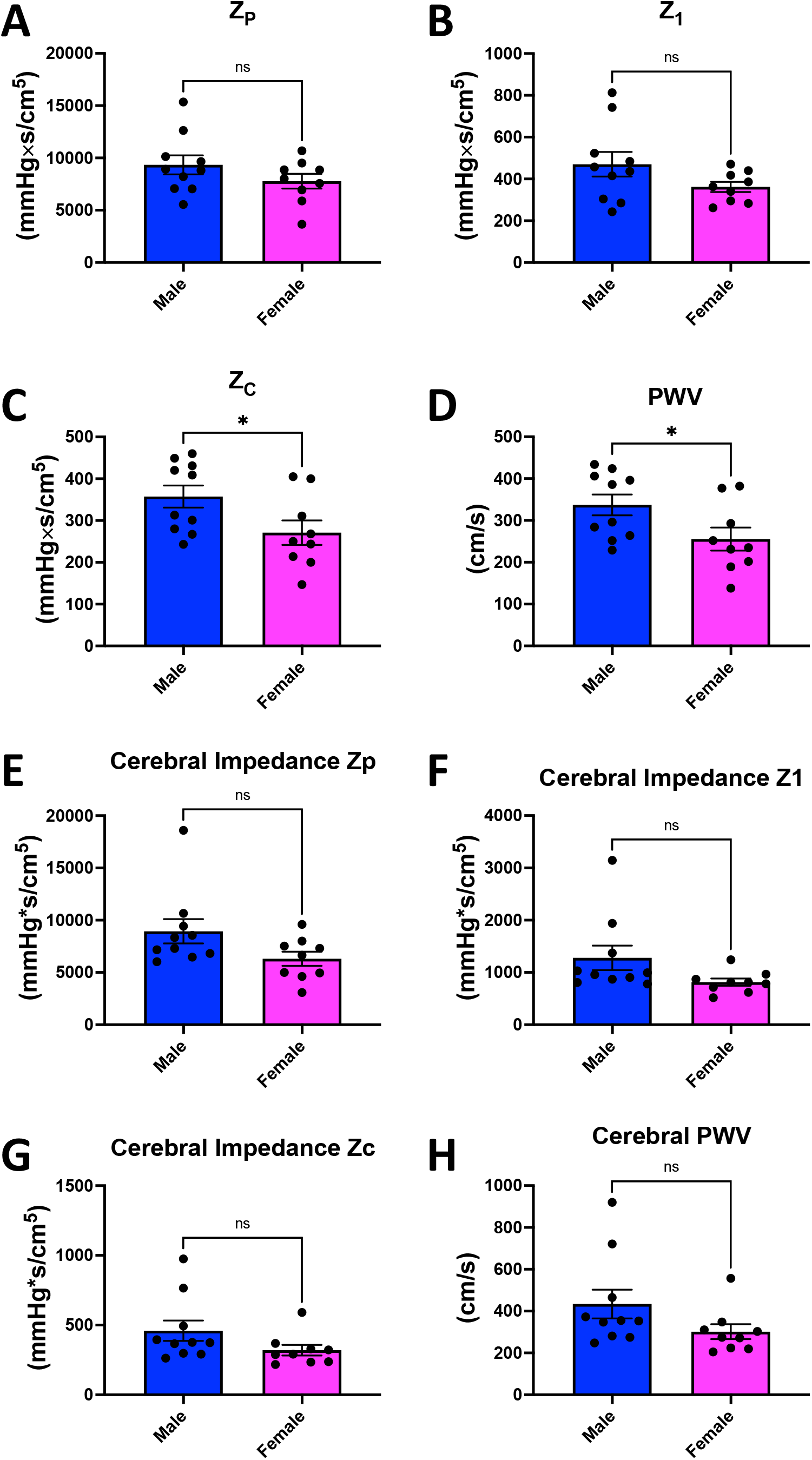
Parameters of aortic and cerebral impedance sex-dependent differences in high fat diet-fed 5-month-old 5XFAD mice. (**A**) Total peripheral resistance (Z_P_), (**B**) impedance at first harmonic (Z_1_), (**C**) characteristic impedance, (Z_C_) (**D**) and impedance-based pulse wave velocity (PWV) in aortic and (**E-H**) cerebral impedance. Data are presented as mean±SEM (n = 9-10/group). * represents p <0.05, ns indicates a statistically non-significant relationship, as determined through an unpaired t-test.

Finally, we sought to apply this method to mice undergoing an acute chronic stress condition. Prior to subjecting 5XFAD mice to stress, non-invasive parameters were collected, as described above. Looking at Tei index, a measure of myocardial performance and cerebral separation distance, another measurement allowed for by non-invasive techniques, we saw no significant differences (**Figure S4A-B**). Additionally, measuring pulse transit time (PTT), which represents the interval for the aortic or cerebral pulse pressure wave to travel from the aortic or cerebral valve to a peripheral site (27), and PWV showed no significant differences for aortic or cerebral measurements (**Figure S4C-D**).

Given that, this non-invasive method validated sex-dependent PWV was minimally different before stress, we subjected a cohort of 5XFAD mice to 1 hour of acute stress. When sexes were grouped together, we saw that every measurement of aortic impedance, Zp, Z1, Zc, or PWVz, was significantly higher in stressed mice (**Figure 5A-D)**. Additionally, cerebral Zp and Z1 were also found to be significantly higher in the cohort subjected to stress, as compared to a non-acutely stressed chow-fed 5XFAD cohort (**Figure 5E-H**). While we investigated sex-dependent differences in the stressed cohort and between cohorts, generally these were non-significant (**Figure S5**). Together, these results validate that our method of aortic and cerebral methods can effectively use Doppler measurements to observe differential chronic diet-dependent and acute stress-induced changes in hemodynamics in 5xFAD and WT mice cohorts.

## Discussion

### Aortic and Cerebral impedance in 5XFAD

When evaluating the aortic peripheral vascular resistance (Zp,), strength of wave reflections from the periphery (Z1), characteristic impedance (Zc,), and PWV we did not note a significant sex-dependent difference in both control and 5XFAD mice compared to their control counterpart (**Figure 3-4**). Notably, in both control and 5XFAD mice we did see that only females displayed increased cerebral impedance. In the case of the WT, females show a slightly lower cerebral impedance Zp, while the 5XFAD model differentially shows decreased Zc and PWV. This can imply that neurovascular coupling or conduction of electrical signals within the brain may be uniquely protective in females with AD. Interestingly, one past study suggested that cerebral blood flow is increased in AD risk states, suggesting a complex relationship between cPWV and AD that must be further investigated (28).

While 5XFAD mice serve as a strong model for human AD that has been used in the past, it lacks neurofibrillary tangles which human models would otherwise have (17). Additionally, while AD in humans has clear sex-dependent differences with increased CVD risk (33, 34), a lesser effect was observed here. This may be due to mouse estrous cycles which we did not consider here (35); thus, it is unclear whether our findings can be extrapolated to humans. Another limitation of our study is that we did not measure Aβ deposition levels, which may be correlated with PWV (30), or other molecular markers of AD pathology. Therefore, it is possible that our findings do not fully capture the complex relationship between hemodynamic parameters and AD pathology. Finally, our study only examined one-time points (12 weeks), and it is possible that longer development of mice may have different effects on hemodynamic parameters and AD pathology, as the latter is linked to aging.

Beyond this, future studies may look at the compounding effects of other experimental changes, such as induced hypertension, as the effectors that modulate sex-dependent differences in aortic and cerebral PWV of 5XFAD mice. Furthermore, there is an unmet need of understanding how aortic and cerebral PWV changes across the aging process for 5XFAD. Aging is especially important to study in the context of 5XFAD as they present a truncated development of AD pathology compared to human counterparts, marked by lower levels of Aβ with isomerized D7 and logarithmic plaque deposition beginning at 2 months of age, indicating that aging can uniquely affect 5XFAD mice (29, 30). Furthermore, past studies have shown that while changes in pulse pulsatility, are more common in younger individuals, stiffness is more likely to arise in older AD patients (31, 32). While 5xFAD females had less severe cardiovascular function impairments than is observed in human AD (33, 34), it is possible that compounding factors such as age and estrous cycles are implicated in our findings (35), highlighting the importance of future studies considering stratifications across sex of PWV while considering these factors. Investigating these effectors may also pave the way for future therapies, as PWV-dependence on estrogen receptors suggests a future avenue of estrogen-driven therapies to mitigate sex-dependent differences in AD.

### Aortic and Cerebral impedance in HFD

While a diet-dependent difference was not observed in 5XFAD, we did not that sex-dependent differences did not follow the same pattern in the chow and HFD cohorts. Notably, characteristic impedance (Zc; average of 2–10 harmonics of impedance modulus), and pulse wave velocity (PWV; aortic stiffness index) were both significantly lower in 5xFAD HFD females mice compared to their male counterparts (**Figure 5**). Both of these measures of aortic impedance are methods of considering the change of pressure in relation to changes in velocity to consider hydraulic external load experienced by the left ventricle (36). Reduced parameters of aortic impedance generally signify a reduced aortic wall stiffness (36). Similarly, PWV typically changes concomitantly with pulse pressure, and high PWV, a marker of arterial stiffness, is known to occur across aging and with AD (13). Even in healthy adults, PWV generally increases with aging (37), so it is possible that this sex-dependent difference may lessen in an older mice cohort. Furthermore, PWV commonly decreases alongside blood pressure in the case of medications such as anti-hypertensives (38). Notably, PWV, which is measured at end-diastolic pressure, can be independent of blood pressure, which is measured at peak systolic pressure, due to less noticeable effects of stiffness in end-diastolic (24). It is still, however, possible that the HFD-induced increase in PWV in male is simply indicative of the higher BP in the 5xFAD HFD cohort. However, given that increased PWV is commonly observed in AD (13), it may also point toward certain cardiovascular differences in the 5xFAD model.

Contrastingly, when looking at cerebral impedance measurements, a sex-dependent difference is no longer observed in the HFD cohort. While variability in the male cohort is high, this is unlikely to account for this difference. Thus, it is possible that at baseline, while 5XFAD females are more resistant to elevated cerebral impedance, chronic diet change ameliorates this advantage. This finding highlights the potential of understanding the confluence of sex and gender. Notably, a prior study showed that while N-acetylneuraminic acid is responsible for HFD-induced increased cognitive impairment in 5xFAD, RNA-sequencing showed diet and AD have different roles in the microglia (39). This suggests that while HFD and AD may affect each other in the brain, they may remain independent effectors in the heart. Other studies have found that HFD modulates plasma metabolites more so than other factors in 5xFAD mice and amplifies the role of sex (40), showing the necessity of exploring sex-dependent differences on diet in the future. Past studies have suggested that female 5xFAD mice, while more susceptible to metabolic dysfunction, HFD may serve a protective mechanism (41).

One other potential avenue in which AD pathology has an interplay with HFD-induced changes are through mitochondria. Previous studies have shown that HFD had depressed respiration concomitant with reduced bioenergetics in the liver (42). In Alzheimer’s Disease, mitochondria are known to decline in axons (43), in 5xFAD mice mitochondria undergo fission in an age-dependent manner which is a driving force between loss of cognitive function and pathology progression observed in 5xFAD mice (44). This loss of function may be partially attributed to oxidative stress with accumulates across aging in 5xFAD mice (45). The role of mitochondria in AD pathology is further underscored by drugs that target voltage-dependent anion channel-1 to protect against pathology by stopping mitochondria dysfunction (46). While these effects have been observed mainly to occur in neurons, future studies may explore if mitochondrial function is also affected in the heart. Notably, in HFD cases alone, mitochondrial oxidative phosphorylation is actually increased which protects against contractile function changes in heart failure (47). This is despite increased oxidative stress, which impairs mitochondrial function, induced by HFD in the brain in mice (48). One consideration is that few studies have yet looked at oxidative stress in cardiac tissue, which may have differential regulation than neuronal oxidative stress (49). Nonetheless, these contrasting roles between HFD and 5XFAD conditions may partially explain some of the altered PWV that we observed.

Future studies may look at the compounding effects of other experimental changes, such as high-salt diet changes in addition to HFD, as well as measuring if there is differential plaque formation under experimental conditions of 5xFAD mice. Beyond this, the specific composition of the diet may strongly affect 5xFAD’s response. For example, while in the past a HFD causes increased neuroinflammation and plaque accumulation (50), a ketogenic diet (high fat/low-carbohydrate) improved cognition through inverse effects of reducing neuroinflammation (51), which may reduce cardiovascular deficits. This may be due to the glycemic content of a diet more heavily influencing the effects on neuroinflammation (52). Thus, this suggests that depending the composition of high-fat relative to salt and carbs can alter positive or negative effects of it; thus, future studies may examine hemodynamic alterations with a high-fat, salt, and sugar diet model (53), which causes increased neuroinflammation in WT mice, to better understand the influence of diet akin to that of the western-populations on AD pathology. Beyond this, since we had middleaged 5xFAD mice (i.e., 4-5 months), it is possible that younger or older mice would respond differently to an HFD due to altered non-linear Aβ accumulation that occurs in 5xFAD mice (29).

### Aortic and Cerebral Impedance in Acute Stress Conditions

Finally, we examined acute stress conditions. Despite baseline 5XFAD conditions before stress exposure, stress was significant different than the 5XFAD mice they were compared to. These results generally showed that all measurements of aortic impedance had a significant increase following stress (**Figure 6**). Additionally, cerebral impedance measures of Zp and Z1 were increased. Given that chronic stress has arisen as a risk factor for AD (54)p, these findings suggest a poorly explicated interplay through which stress may contribute to AD pathology through cerebrovascular effects.

**Figure 6:**
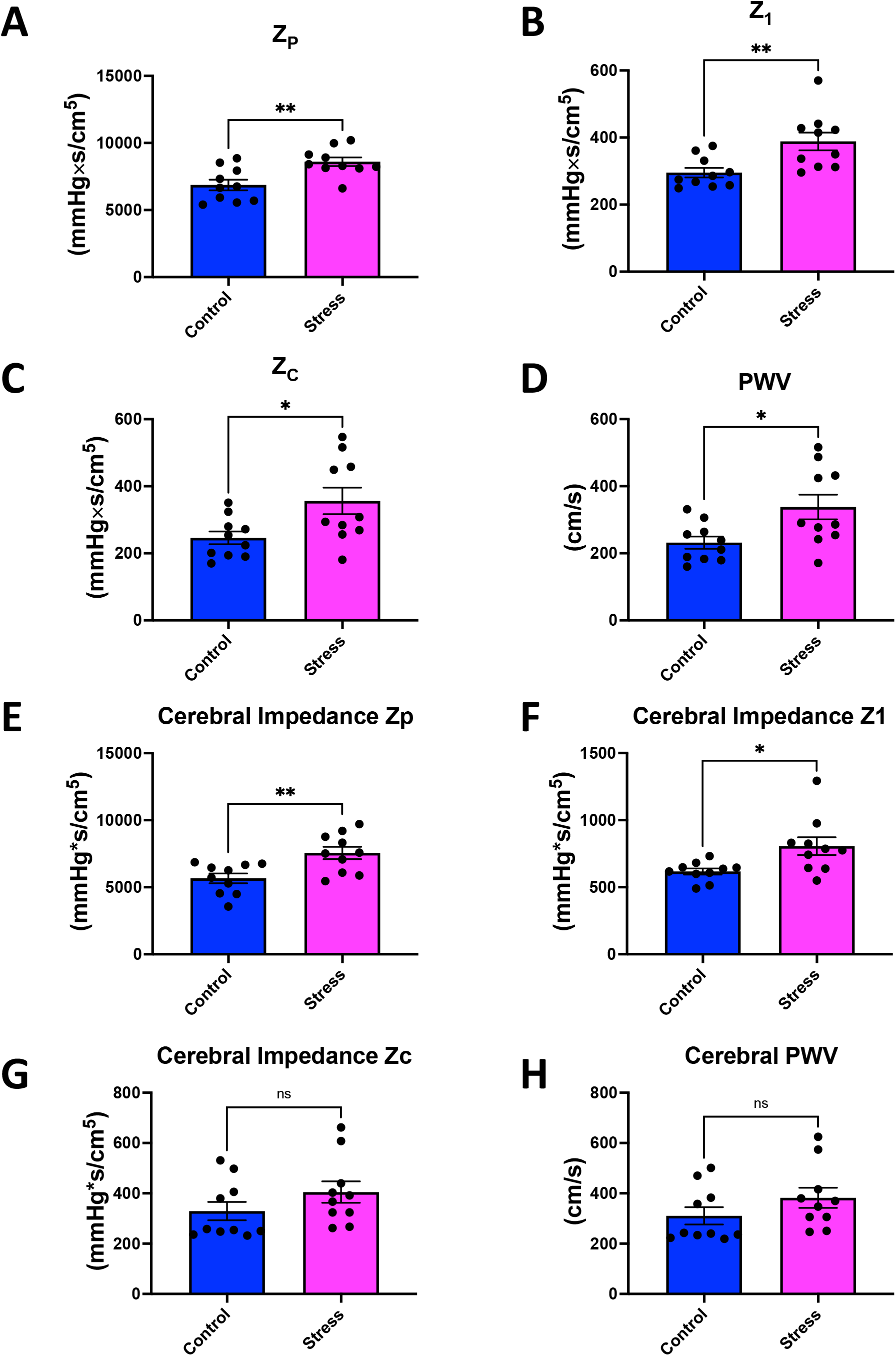
Parameters of aortic and cerebral in pre-stress and stressed state of 5-month-old 5XFAD mice. (**A**) Total peripheral resistance (Z_P_), (**B**) impedance at first harmonic (Z_1_), (**C**) characteristic impedance, (Z_C_) (**D**) and impedance-based pulse wave velocity (PWV) in aortic and (**E-H**) cerebral impedance. Data are presented as mean±SEM (n = 10/group). *, ** represents p <0.05, p<0.01, respectively, and ns indicates a statistically non-significant relationship, as determined through an unpaired t-test.

Notably, sex-dependent differences were not drastic in cerebral impedance of the HFD cohort (**Figure S5**). This suggests that, like an HFD challenge, while females may have slightly lower cerebral impedance at baseline in 5XFAD, these sex-dependent differences may fade under certain chronic or acute challenges. These findings suggest that the relationship between PWV and AD pathology in animal models may be more complex than previously thought and highlight the importance of studying multiple hemodynamic parameters in the context of AD. Females may be less vulnerable to AD partially due to reduced cerebral impedance Zp, however global increases in aortic and cerebral PWV following stress induction may erase sex-dependent differences. Interestingly, past studies have shown that stress in females only, and not males, leads to elevated beta-amyloid (55). This highlights that PWV and cPWV are distinct from overall beta-amyloid accumulation, and not necessarily correlative, emphasizing the importance of continuing to study PWV and cPWV in a range of conditions.

Further studies are needed to elucidate the mechanisms underlying these findings and their implications for the development and progression of AD. In conclusion, our study showed no significant changes in most of the sex-dependent hemodynamic parameters examined, with the most pertinent change but confirmed the deleterious effects of stress, which displayed slight sex-dependent differences.

## Conclusions

5XFAD mice are important new experimental models for the study of Alzheimer’s Disease (17), but such models have limited research evaluating their *in situ* global cardiovascular function. Together our data also highlight that AD pathology should not only be considered in the context of tau pathology and cognitive decline but aortic and cerebral impedance metrics as well. In example, recent studies have demonstrated that Resveratrol can reduce HFD-induced accelerated cognitive decline in 5xFAD through proteolytic mechanisms (56), but these same pathways may not change HFD-induced cardiovascular remodeling. Thus, the impact of AD on cardiovascular parameters, especially under altered environmental states must be considered. Beyond using noninvasive measurements of Doppler mitral inflow and aortic flow velocity, invasive measurements of aortic and LV pressure, and the calculated aortic impedance, we also evaluated cerebral impedance.

In the future, cPWV may also aid in the early detection and monitoring of other neurological disorders. While arterial PWV may serve as a mechanism to explore cerebral perfusion, cerebral microbleeds have arisen as markers that are important to study in AD (8). Other studies have also shown that glutamate chemical exchange saturation transfer has been observed to concomitantly decrease with cerebral blood flow in 5XFAD mice, suggesting other novel imaging techniques that can potentially be used alongside cPWV (57). Another promising technique is 4D flow MRI, which has been used to show increased transcranial PWV in AD (58). Therefore, further studies are needed to elucidate the mechanisms underlying how cPWV and other innovative techniques may explore AD in 5XFAD and other models.

### Limitations

#### Study

There are several limitations to this study that must be considered when interpreting the results. While 5xFAD mice have many similarities to AD pathology in humans, it lacks neurofibrillary tangles and may have altered cardiac pathways which are poorly elucidated. Different mouse models of AD also respond to HFD differently (59, 60), while it remains unclear which most closely mimics that of humans which limits the ability to extrapolate findings to AD (17, 61). Beyond this, while past studies have clearly shown HFD-induced accelerated cognitive decline in 5xFAD mice (56), we did not specifically look at this or correlate brain hemodynamics or other neuronal molecular markers of AD pathology with cardiovascular stiffness changes. While we sought more to understand the impact of 5xFAD condition on diet-induced changes, it is possible there are additional neglected links between hemodynamics in the opposite regulation of AD. While we began an HFD at 5 months of age, the age mice can affect response to HFD, with past studies showing that HFD feeding before 3 months of age can have protective effects against cognitive decline (62). Past studies also observed high- and low-weight cohort differences in response to HFD in 5xFAD, which we did not see, is exacerbated in female mice (62).

#### Methods

Generally, these models are highly reproducible and adapted from existing aortic PWV measurements, so they require minimal new equipment. However, an important limitation is that wall motion must be converted to wave form in mice.

## FUNDING

This work is supported by National Institute of Health (NIH) NIDDK T-32, number DK007563 entitled Multidisciplinary Training in Molecular Endocrinology to Z.V.; NSF MCB #2011577I to S.A.M.; The UNCF/Bristol-Myers Squibb E.E. Just Faculty Fund, Career Award at the Scientific Interface (CASI Award) from Burroughs Welcome Fund (BWF) ID # 1021868.01, BWF Ad-hoc Award, NIH Small Research Pilot Subaward to 5R25HL106365-12 from the National Institutes of Health PRIDE Program, DK020593, Vanderbilt Diabetes and Research Training Center for DRTC Alzheimer’s Disease Pilot & Feasibility Program. CZI Science Diversity Leadership grant number 2022-253529 from the Chan Zuckerberg Initiative DAF, an advised fund of Silicon Valley Community Foundation to A.H.J.; National Institutes of Health grants R01HL147818 and R01HL144941 (A. Kirabo). Its contents are solely the responsibility of the authors and do not necessarily represent the official view of the NIH. The funders had no role in study design, data collection and analysis, decision to publish, or preparation of the manuscript.

### Disclosures

Dr. Reddy is a collaborator and consultant with Indus Instruments, Webster, TX. All other authors have no competing interests.

### Author Contributions

A–M - 3456, –N - 3456, –V - 3456, H–B - 34, –G - 34, –V - 34, –B - 34, –E - 34, –A - 34, AM – 34, ES-34, AC-34, JD -34, DS – 34, SD – 34, T–P - 23, J–G - 678, –E - 678, –D - 678, M–E - 7,8 G–T - 5678, A–H - 5678, A–R - 12345678

1 Conceived and designed research, 2 Performed experiments and Data collection, 3 Data analysis, 4 Prepared figures, 5 Interpretation of results, 6 Drafted manuscript, 7 Edited and revised manuscript, 8 Approved final version of manuscript.

## Supplementary Files

**Supplementary Figure 1:** Workflow of inducing a high-fat diet or stress experimental conditions in 5XFAD mice.

**Supplementary Figure 2:** Flow of Doppler, electrocardiography (ECG), and blood pressure (BP) signals from the animal to the transceiver and amplifier to generate the Doppler inphase (I) and quadrature (Q) audio signals, ECG, and BP signals. The Doppler Flow Velocity System (DFVS) hardware consists of a high-speed digitizer which sends the digitized I/Q, ECG, & BP to the DFVS software for acquisition, display, storage, and analysis.

**Supplementary Figure 3:** Parameters of aortic and cerebral impedance in a mixture of male and female 5XFAD mice on chow and high-fat diets. (**A**) Total peripheral resistance (Z_P_), (**B**) impedance at first harmonic (Z_1_), (**C**) characteristic impedance, (Z_C_) (**D**) and impedance-based pulse wave velocity (PWV) in aortic and (**E-H**) cerebral impedance. Data are presented as mean±SEM (n = 19/group). ns indicates a statistically non-significant relationship, as determined through an unpaired t-test.

**Supplementary Figure 4:** Parameters of aortic and cerebral impedance in 5XFAD mice measured through noninvasive methods in males and females. (**A**) Myocardial performance index (Tei index), a measure of the overall function of the heart, which takes into account systolic and diastolic function, as calculated by (IVCT+IVRT)/ET. (**B**) Cerebral separation distance. (**C**) Pulse transit time and (**D**) impedance-based pulse wave velocity (PWV) in aortic and (**E-F**) cerebral parameters. Data are presented as mean±SEM (n = 5/group). ns indicates a statistically non-significant relationship, as determined through an unpaired t-test.

**Supplementary Figure 5:** Parameters of aortic and cerebral impedance sex-dependent differences in non-stressed and stressed 5XFAD mice. (**A**) Total peripheral resistance (Z_P_), (**B**) impedance at first harmonic (Z_1_), (**C**) characteristic impedance, (Z_C_) (**D**) and impedance-based pulse wave velocity (PWV) in aortic and (**E-H**) cerebral impedance. Data are presented as mean±SEM (n = 5/group). * represents p <0.05, ns indicates a statistically non-significant relationship, as determined through an unpaired t-test.

**Supplementary Table 1:** Summary of Pulse Wave and Cerebral Impedance Measurements Utilized.

