## Supplement for "Methods to Utilize Pulse Wave Velocity to Measure Alterations in Cerebral and Cardiovascular Parameters"

### Supplementary Figure

Figure S1

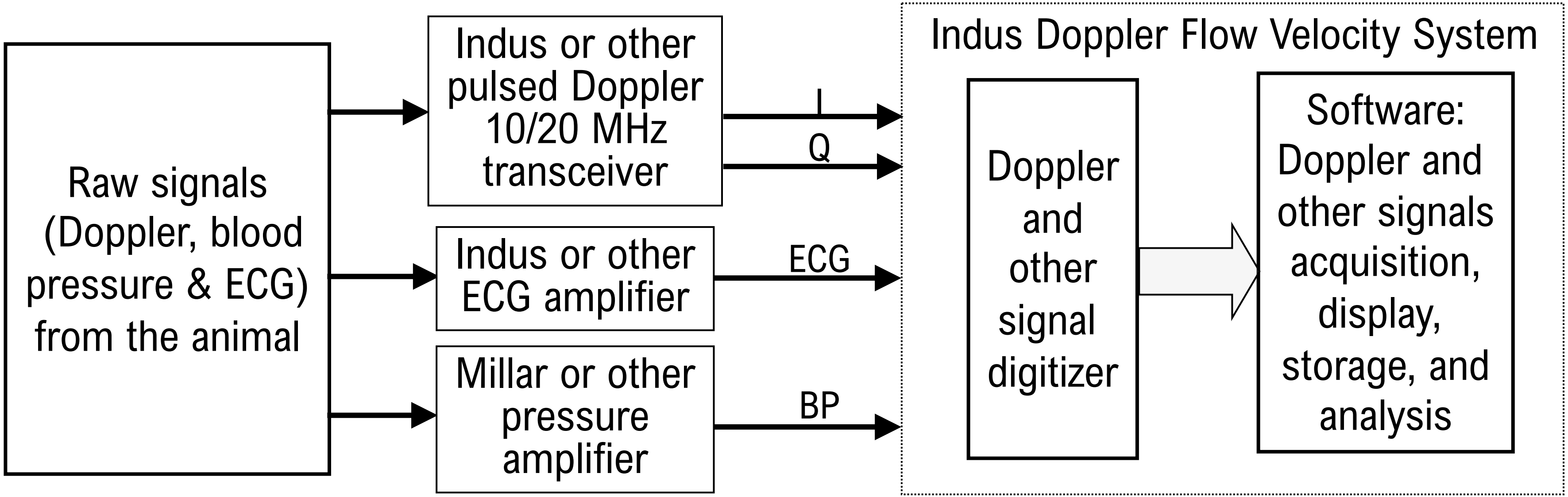
Raw signals (Doppler, blood pressure & ECG) from the animal
Indus or other pulsed Doppler 10/20 MHz transceiver
Indus or other ECG amplifier
Millar or other pressure amplifier
I
Q
ECG
BP
Indus Doppler Flow Velocity System
Doppler and other signal digitizer
Software: Doppler and other signals acquisition, display, storage, and analysis

Figure S2

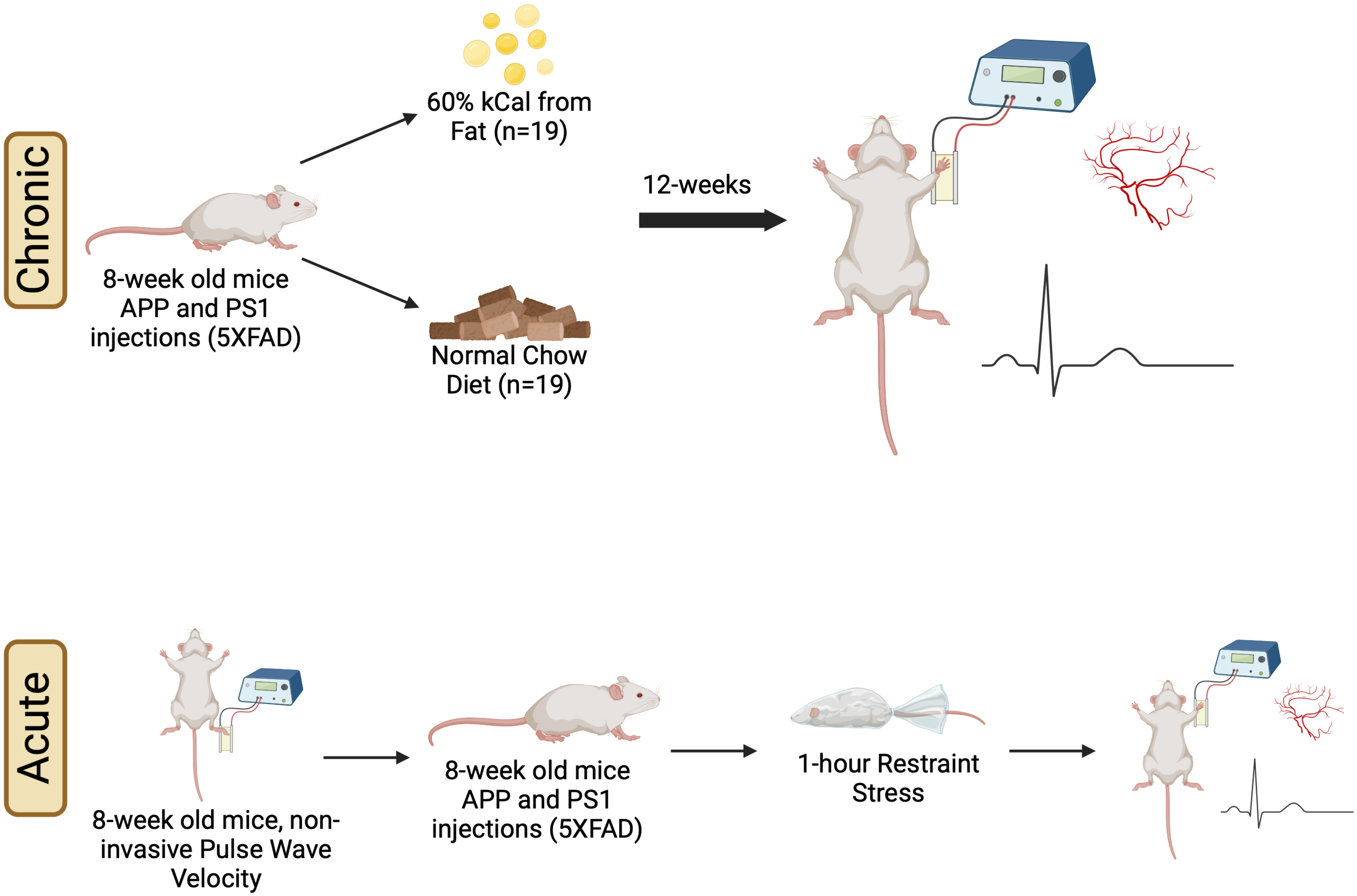
Chronic
60% kCal from Fat (n=19)
12-weeks
8-week old mice APP and PS1 injections (5XFAD)
Normal Chow Diet (n=19)
Acute
8-week old mice, non-invasive Pulse Wave Velocity
8-week old mice APP and PS1 injections (5XFAD)
1-hour Restraint Stress

Figure S3

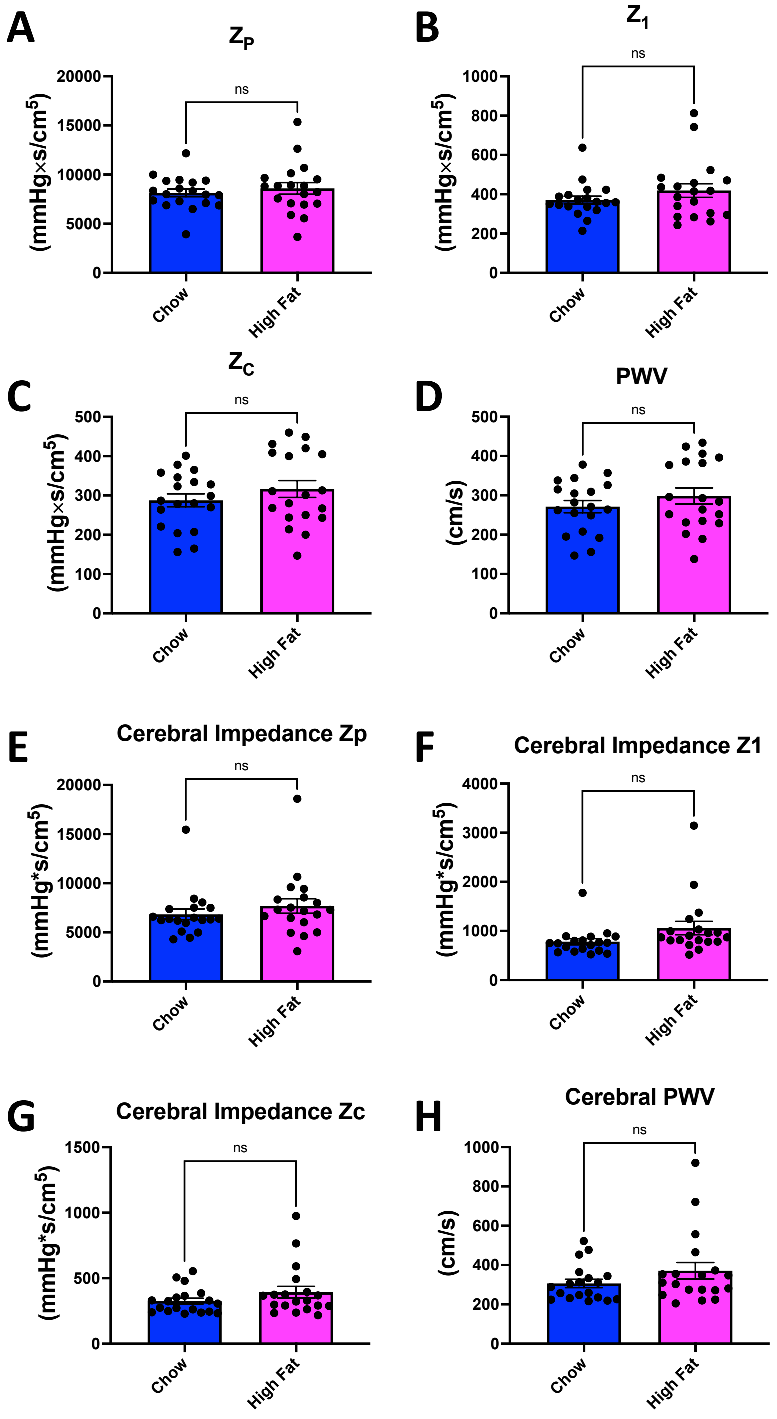
A
Z_P
ns
20000
15000
10000
5000
0
(mmHg×s/cm5)
Chow
High Fat
B
Z_1
ns
1000
800
600
400
200
0
(mmHg×s/cm5)
Chow
High Fat
C
Z_C
ns
500
400
300
200
100
0
(mmHg×s/cm5)
Chow
High Fat
D
PWV
ns
500
400
300
200
100
0
(cm/s)
Chow
High Fat
E
Cerebral Impedance Zp
ns
20000
15000
10000
5000
0
(mmHg*s/cm5)
Chow
High Fat
F
Cerebral Impedance Z1
ns
4000
3000
2000
1000
0
(mmHg*s/cm5)
Chow
High Fat
G
Cerebral Impedance Zc
ns
1500
1000
500
0
(mmHg*s/cm5)
Chow
High Fat
H
Cerebral PWV
ns
1000
800
600
400
200
0
(cm/s)
Chow
High Fat

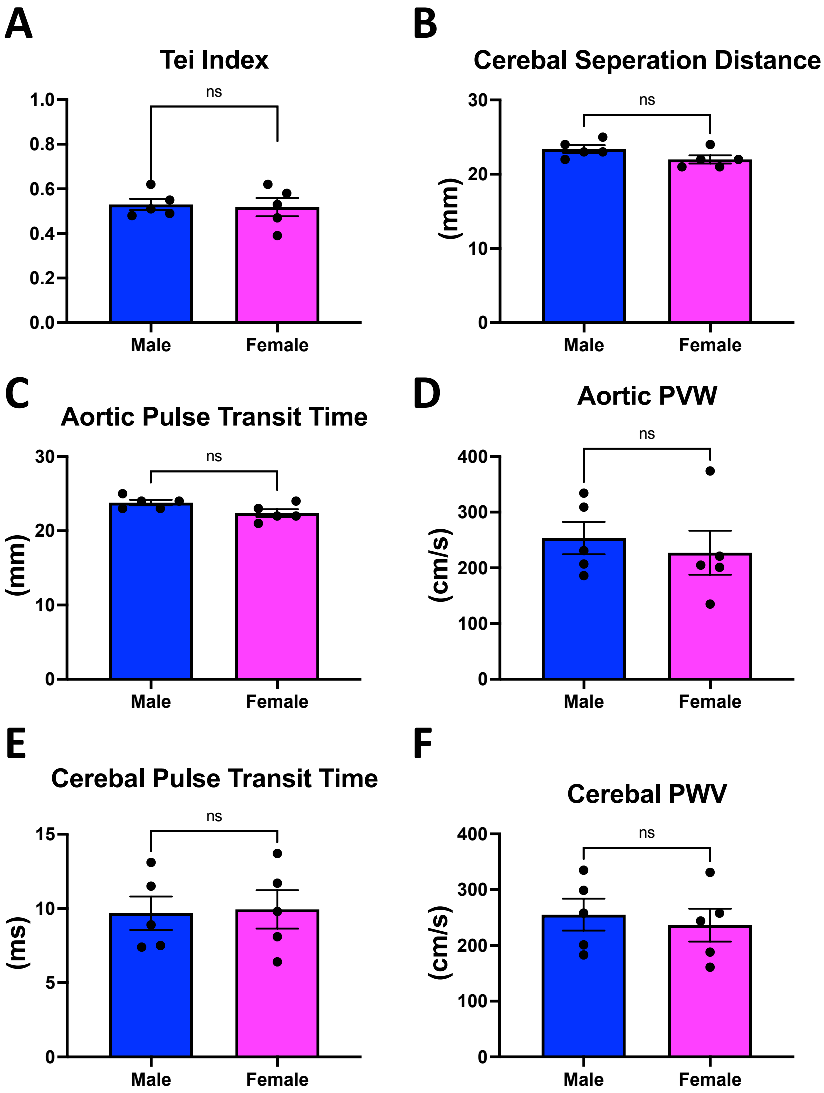
A
Tei Index
ns
1.0
0.8
0.6
0.4
0.2
0.0
Male
Female
B
Cerebal Seperation Distance
ns
30
20
10
0
(mm)
Male
Female
C
Aortic Pulse Transit Time
ns
30
20
10
0
(mm)
Male
Female
D
Aortic PVW
ns
400
300
200
100
0
(cm/s)
Male
Female
E
Cerebal Pulse Transit Time
ns
15
10
5
0
(ms)
Male
Female
F
Cerebal PWV
ns
400
300
200
100
0
(cm/s)
Male
Female

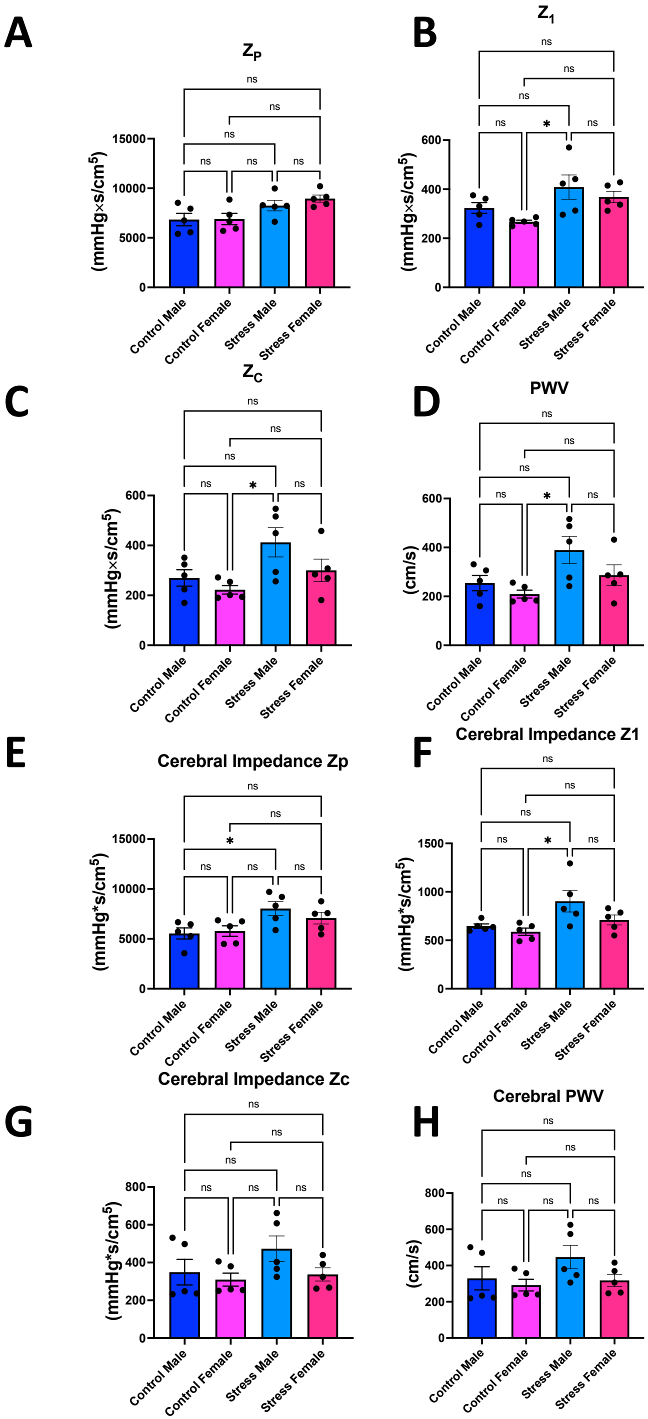
A
Zp
ns
ns
ns
ns
ns
ns
15000
10000
5000
0
(mmHg×s/cm5)
Control Male
Control Female
Stress Male
Stress Female
B
Z1
ns
ns
ns
ns
*
ns
600
400
200
0
(mmHg×s/cm5)
Control Male
Control Female
Stress Male
Stress Female
C
Zc
ns
ns
ns
ns
*
ns
600
400
200
0
(mmHg×s/cm5)
Control Male
Control Female
Stress Male
Stress Female
D
PWV
ns
ns
ns
ns
*
ns
600
400
200
0
(cm/s)
Control Male
Control Female
Stress Male
Stress Female
E
Cerebral Impedance Zp
ns
ns
*
ns
ns
ns
15000
10000
5000
0
(mmHg*s/cm5)
Control Male
Control Female
Stress Male
Stress Female
F
Cerebral Impedance Z1
ns
ns
ns
ns
*
ns
1500
1000
500
0
(mmHg*s/cm5)
Control Male
Control Female
Stress Male
Stress Female
G
Cerebral Impedance Zc
ns
ns
ns
ns
ns
ns
800
600
400
200
0
(mmHg*s/cm5)
Control Male
Control Female
Stress Male
Stress Female
H
Cerebral PWV
ns
ns
ns
ns
ns
ns
800
600
400
200
0
(cm/s)
Control Male
Control Female
Stress Male
Stress Female

| Measurement | Definition |
|---|---|
| **Impedance at first harmonic (Z1)** | Blood flow resistance at the frequency of the first harmonic of the pulse wave, which is an indicator of arterial walls stiffness. |
| **Characteristic impedance (ZC)** | The characteristic impedance of the pulse wave propagation without reflection. In order words, impedance at zero frequency, which offers information about arterial compliance and wave reflection characteristics. |
| **Total peripheral resistance (ZP)** | Resistance to blood flow in the systemic circulation, as measured by combined effects of factors including blood vessel diameter and blood viscosity. Indicator of overall cardiovascular health and represents the average 2nd to 10th harmonic. |
| **Impedance-based pulse wave velocity (PWV)** | Measure of the velocity of the arterial pulse wave as it travels through the arterial tree. |
| **Cerebral Z1** | Resistance to blood flow at the frequency of the first harmonic in the cerebral circulation, which is an indicator of cerebral arterial stiffness. |
| **Cerebral ZC** | The characteristic impedance of the cerebral pulse wave propagation with reflection, reflects the properties of the cerebral arteries. |
| **Cerebral ZP** | Measure of the resistance to blood flow in the cerebral circulation which is an indicator of the state of the cerebral vasculature. |
| **Impedance-based cerebral PWV** | Speed at which the pulse wave travels through the cerebral arteries, as measurement of cerebral arterial stiffness. |
